# MakeMyFigure: An Interactive Platform for Reproducible Quantitative Data Visualization, Analysis, and Scientific Figure Construction

**DOI:** 10.64898/2026.09.15.751916

**Authors:** Suresh Poudel, Him K. Shrestha, Jeremy C. Crawford, Fabio Demontis, Douglas R. Green

## Abstract

Reproducible data analysis and visualization are essential for reliable biological and biomedical research, yet creating high-quality, publication-ready figures often requires moving data and results between multiple analysis, visualization, and graphics tools. This fragmented, multi-layered process can make it difficult to trace how individual figure panels were generated, preserve the underlying analytical decisions, and reproduce them later. Here, we introduce MakeMyFigure, a free and open-source platform for data visualization, analysis, and creation of multi-panel scientific figures. MakeMyFigure integrates data processing, statistical analysis, visualization, and figure assembly within a single workflow. It supports a range of quantitative data formats and structures, including feature-by-sample matrices and precomputed statistical results, and provides data-aware visualization recommendations to help users select appropriate plots from a library of 38 visualization types. These capabilities allow researchers to move directly from experimental measurements to commonly used statistical analyses and graphical representations without requiring programming expertise. Eighteen statistical procedures are implemented, and their results are drawn directly onto the plot families that support statistical annotation, keeping analytical results linked to the panels they generate. Importantly, MakeMyFigure uses machine-readable JSON specifications to record the identity and checksum of the source data, the processing steps, visualization settings, and statistical parameters used to generate each figure panel. It can also save a figure as a portable package, freezing the specifications together with the data. This allows figures to be regenerated from their recorded specifications and source data, or from the package alone on another computer, rather than relying on manually reconstructed workflows. Using published datasets from several biological fields, independent statistical validation in R, and a comparison with 15 representative analysis, visualization, and figure-generation tools, we show that MakeMyFigure combines accessible, code-free figure creation with panel-level computational reproducibility. Overall, MakeMyFigure provides a unified approach for creating, documenting, and reproducing scientific figures. Source code and documentation are available at https://github.com/surPoudel/make-my-figure, https://github.com/surPoudel/make-my-figure/releases/tag/v1.1.0.

## Introduction

Quantitative biological research increasingly depends on workflows that combine data processing, statistical analysis, visualization, and figure preparation^1^. Although these steps are closely related, they are often performed in separate software environments, requiring researchers to transfer data and results across statistical programs, visualization tools, and graphics editors^2^. Such fragmented workflows can introduce inconsistencies, obscure how analyses were performed, and make it difficult to reproduce how individual figure panels were generated^3–6^. At the same time, many statistical computing environments require programming expertise, creating a barrier for experimental researchers who need to perform standard statistical analyses but do not routinely write code^1,7^. Conversely, dedicated graphics and figure-design software can provide extensive control over visual presentation but typically does not perform statistical analysis or maintain a direct connection to the underlying analytical workflow.

A range of established tools partially bridges these divisions across the analysis-to-figure workflow for biological researchers. General-purpose scientific applications such as GraphPad Prism^8^, OriginPro^9^, JMP^10^, SigmaPlot^11^ and DataGraph^12^ combine interactive data analysis with publication-oriented visualization, whereas jamovi^13^ and JASP^14^ emphasize accessible, code-free statistical analysis. In biological research, platforms such as Perseus^15^, iDEP^16^, ExpressAnalyst^17^, and Galaxy^18^ provide domain-specific workflows for processing, statistical analysis, and visualization, while SRplot^19^ and Hiplot^20^ offer broad collections of web-based scientific and biological visualizations. KNIME^21^ supports reproducible, node-based analytical workflows across heterogeneous data sources, and BioRender^22^ combines scientific graphics, graphing and figure composition within a design-oriented environment. Collectively, these tools provide substantial capabilities for statistical analysis, biological data exploration, visualization, workflow reproducibility, or figure preparation (Table 1), but they generally emphasize a particular stage of the workflow rather than maintaining a unified connection between source data, analytical operations, visualization parameters, and the individual panels of a composite scientific figure.

**Table 1.**
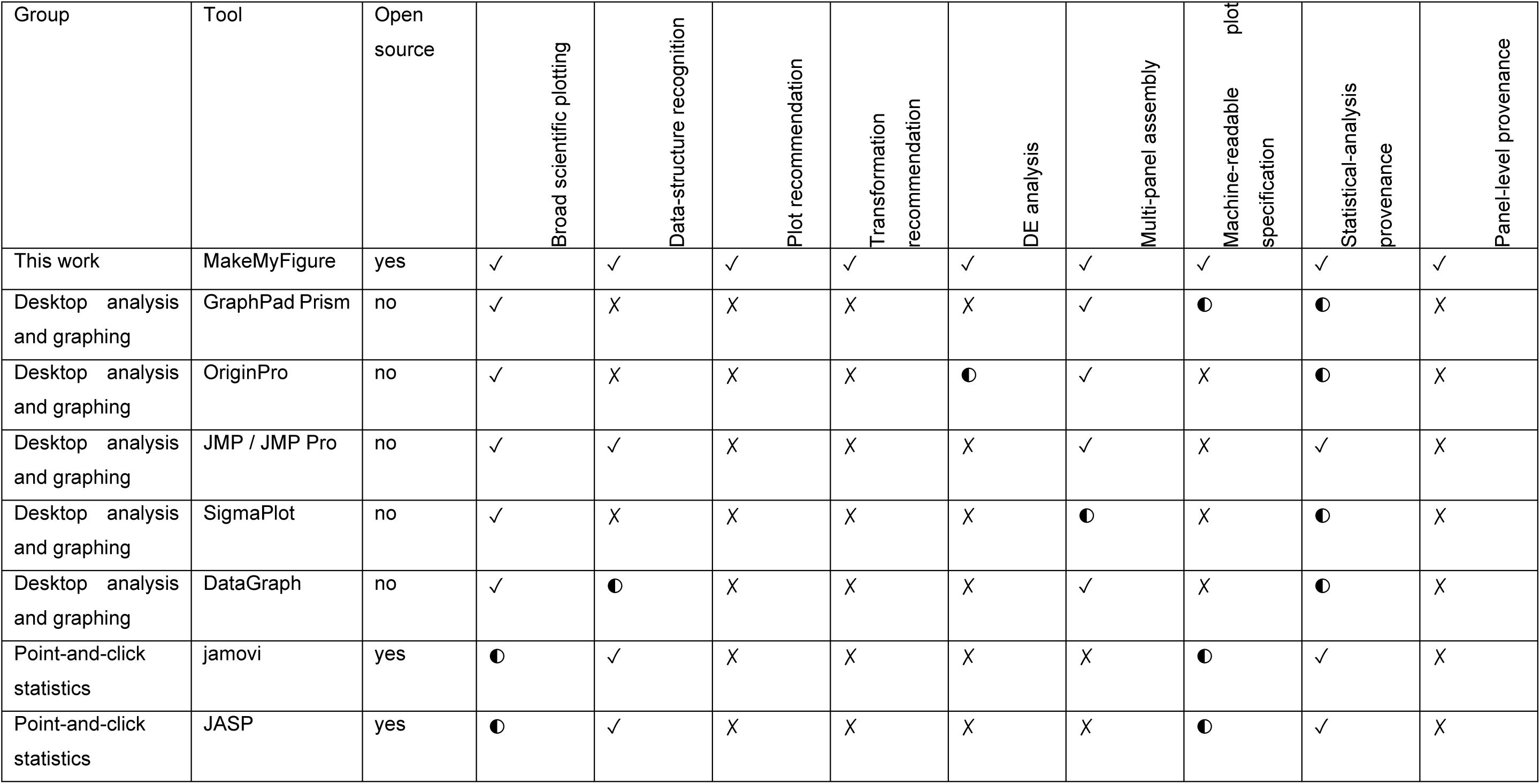
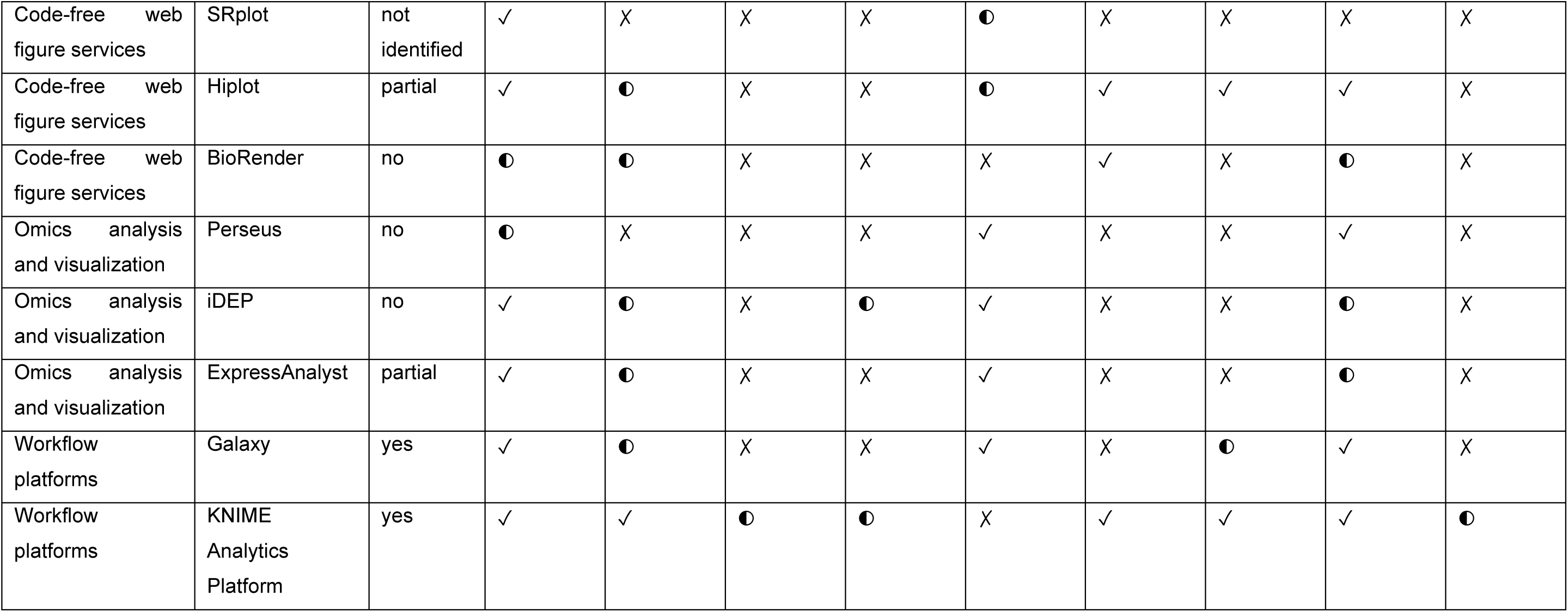
Comparison of MakeMyFigure with representative scientific analysis, visualization and figure-generation tools. Tools are grouped by their primary use case: desktop analysis and graphing, point-and-click statistics, code-free web figure services, omics analysis and isualization, and workflow platforms. Features were assessed based on whether each tool supports broad scientific plotting, automated ecognition of data structure or type, plot recommendation, transformation recommendation, differential expression (DE) analysis, multi-panel igure assembly, export or storage of plot parameters in a machine-readable specification, preservation of statistical analysis provenance, and panel-level provenance. ✓, supported; ◐, partially supported or available with limitations; ✗, not supported or not identified as a native capability. The capabilities of the 15 comparators were assessed using official documentation, peer-reviewed software papers, license files, and source ode. Open source: yes (OSI license), partial (part of the stack open), no (proprietary), or not identified. For MakeMyFigure, DE analysis denotes ts native feature-level differential summary (Welch’s or Student’s t-test, Mann–Whitney U, paired tests, one-way ANOVA or Kruskal–Wallis with multiple-testing correction, reporting log₂ fold changes and effect sizes) on a normalized matrix; a negative-binomial count model (PyDESeq2) s available as an optional extra, and precomputed DE tables from count-model tools are plotted as given.

| Group | Tool | Open source | Broad scientific plotting | Data-structure recognition | Plot recommendation | Transformation recommendation | DE analysis | Multi-panel assembly | Machine-readable plot specification | Statistical-analysis provenance | Panel-level provenance |
| --- | --- | --- | --- | --- | --- | --- | --- | --- | --- | --- | --- |
| This work | MakeMyFigure | yes | ✓ | ✓ | ✓ | ✓ | ✓ | ✓ | ✓ | ✓ | ✓ |
| Desktop analysis and graphing | GraphPad Prism | no | ✓ | X | X | X | X | ✓ | ● | ● | X |
| Desktop analysis and graphing | OriginPro | no | ✓ | X | X | X | ● | ✓ | X | ● | X |
| Desktop analysis and graphing | JMP / JMP Pro | no | ✓ | ✓ | X | X | X | ✓ | X | ✓ | X |
| Desktop analysis and graphing | SigmaPlot | no | ✓ | X | X | X | X | ● | X | ● | X |
| Desktop analysis and graphing | DataGraph | no | ✓ | ● | X | X | X | ✓ | X | ● | X |
| Point-and-click statistics | jamovi | yes | ● | ✓ | X | X | X | X | ● | ✓ | X |
| Point-and-click statistics | JASP | yes | ● | ✓ | X | X | X | X | ● | ✓ | X |
| Code-free web figure services | SRplot | not identified | ✓ | X | X | X | ● | X | X | X | X |
| Code-free web figure services | Hiplot | partial | ✓ | ● | X | X | ● | ✓ | ✓ | ✓ | X |
| Code-free web figure services | BioRender | no | ● | ● | X | X | X | ✓ | X | ● | X |
| Omics analysis and visualization | Perseus | no | ● | X | X | X | ✓ | X | X | ✓ | X |
| Omics analysis and visualization | iDEP | no | ✓ | ● | X | ● | ✓ | X | X | ● | X |
| Omics analysis and visualization | ExpressAnalyst | partial | ✓ | ● | X | X | ✓ | X | X | ● | X |
| Workflow platforms | Galaxy | yes | ✓ | ● | X | X | ✓ | X | ● | ✓ | X |
| Workflow platforms | KNIME Analytics Platform | yes | ✓ | ✓ | ● | ● | X | ✓ | ✓ | ✓ | ● |

Despite these advances, the analytical and graphical stages of figure generation often remain disconnected. Data may be processed and statistically tested in one environment, visualized in another, and subsequently assembled with other panels in a separate figure-design application. Although individual tools may preserve their own analysis state, the resulting composite figure does not necessarily retain a unified, machine-readable record linking each panel to its source data, processing steps, statistical parameters, and visualization settings (Table 1). This limitation becomes particularly important when heterogeneous outputs, such as statistical plots, heatmaps, survival analyses, and images, are combined within a single figure. Thus, while existing platforms provide substantial capabilities for analysis, visualization, workflow reproducibility, or figure composition, integrated support for these functions together with panel-level provenance, a record of how each panel was generated, remains limited.

Here, we present MakeMyFigure, a free and open-source platform for scientific data analysis, visualization, and multi-panel figure construction. MakeMyFigure integrates data processing, statistical testing, visualization, and figure assembly into a single code-free workflow, while recording the identity of the source data, the analytical operations, visualization settings, and statistical parameters for each panel in machine-readable specifications that can be frozen together with the data in a portable figure package. We evaluate MakeMyFigure through the recreation of six published figure panels from their source data, validation of its statistical implementations against independent calculations in R, plotting of published differential-expression and survival results from openly available datasets, reconstruction and reuse of figures from their recorded specifications and portable packages, and tests of its behavior across input formats, computing environments, export formats, data sizes, and repeated rendering, together with a documentation-based comparison with 15 representative tools (Table 1).

## Results

### MakeMyFigure: An application that connects data, analysis, visualization, and figure assembly

A figure generated for the publication of a manuscript usually goes through multiple stages, starting from data to figure rendering (Fig. 1A). Very often, multiple tools are used independently to refine the figure. There is no consistency in measurement and dimensions as measurements are organized in one program, processed and tested in another, plotted and annotated in a third, and assembled with other panels in a graphics editor. Each program serves its own stage, but the context accompanying a panel is not carried across handoffs: the identity of the source table, the processing and test parameters, the plot settings and annotations, and the panel’s position in the composite. Once the figure exists, this context must be reconstructed by hand.

**Figure 1.**
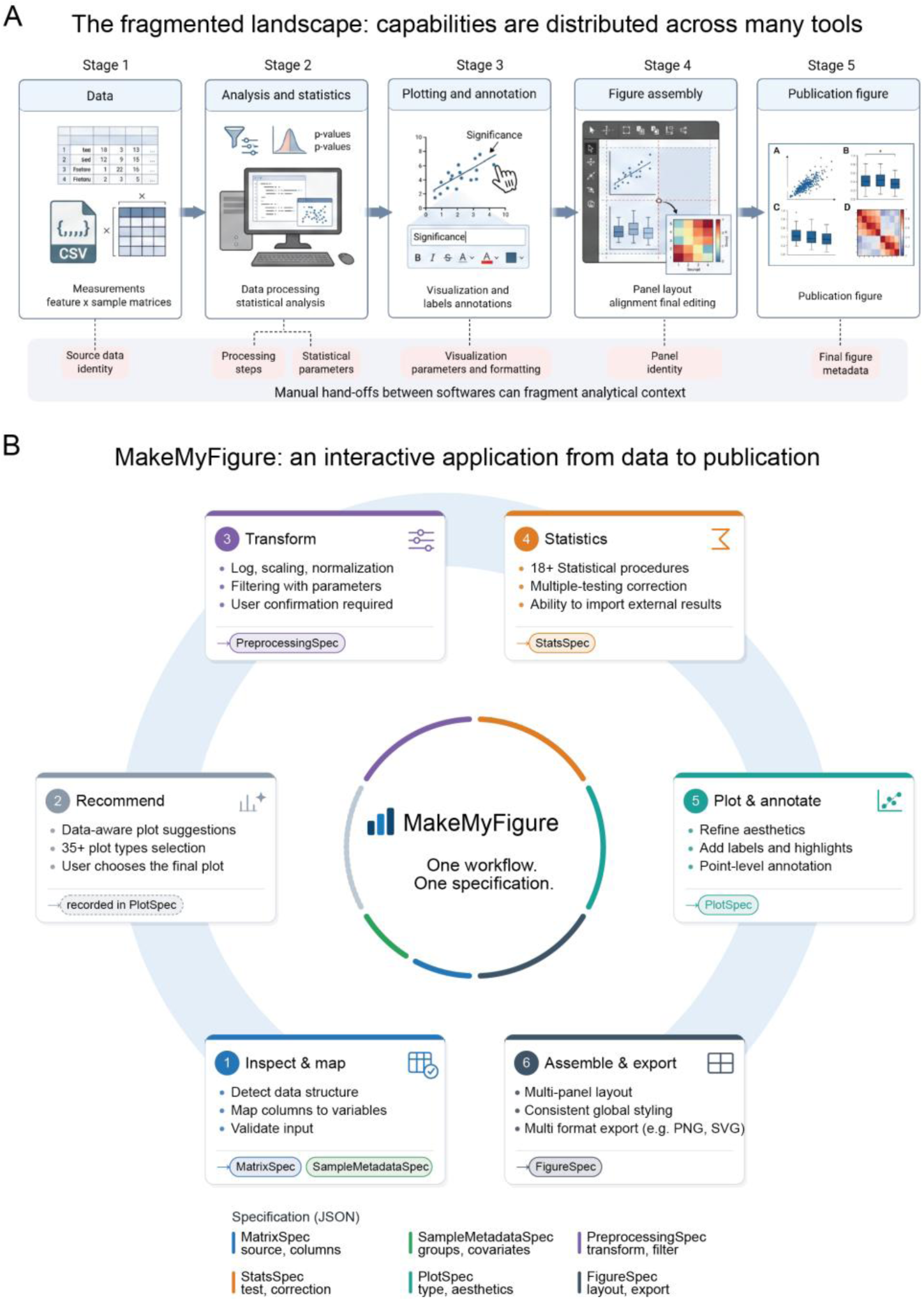
Figure preparation across independent tools versus the connected workflow in MakeMyFigure. (A) Conventional figure preparation spans multiple tools for data preparation, analysis, visualization, and assembly, with contextual information often lost between stages. (B) MakeMyFigure connects these steps into six stages: inspect and map, recommend, transform, statistics, plot and annotate, and assemble and export. Each stage generates a machine-readable specification that can be saved with the data as a reproducible figure package.

MakeMyFigure organizes the same operations into six connected stages within a single application (Fig. 1B). A table or workbook is read, its structure is detected, and column roles are proposed and then confirmed by the user. From the confirmed structure, a rule-based engine proposes candidate plots together with compatible transformations and statistical tests, and the user selects among them. Transformations such as log transformation, filtering, scaling, and normalization are offered for feature-by-sample matrices, with their parameters shown. A transformation is applied only after confirmation and only to a derived copy, so the original table is retained. The route for a feature-by-sample matrix, from confirmed column roles and groups through profiling, suggested preprocessing and a recorded derived matrix to plots, is shown on a deposited count matrix in Fig. 2. Eighteen statistical procedures are registered, with multiple-testing correction across pairwise comparisons; differential expression is computed natively for feature-by-sample matrices, and precomputed results, such as differential-expression tables from other tools, are plotted as given rather than recalculated. Plots are drawn from a registry of 38 plot types via a single styling layer, refined with labels, highlights, and point-level annotations, and assembled into a multi-panel figure with consistent styling, then exported to raster and vector formats. A desktop application and a local browser application call the same core implementation.

**Figure 2.**
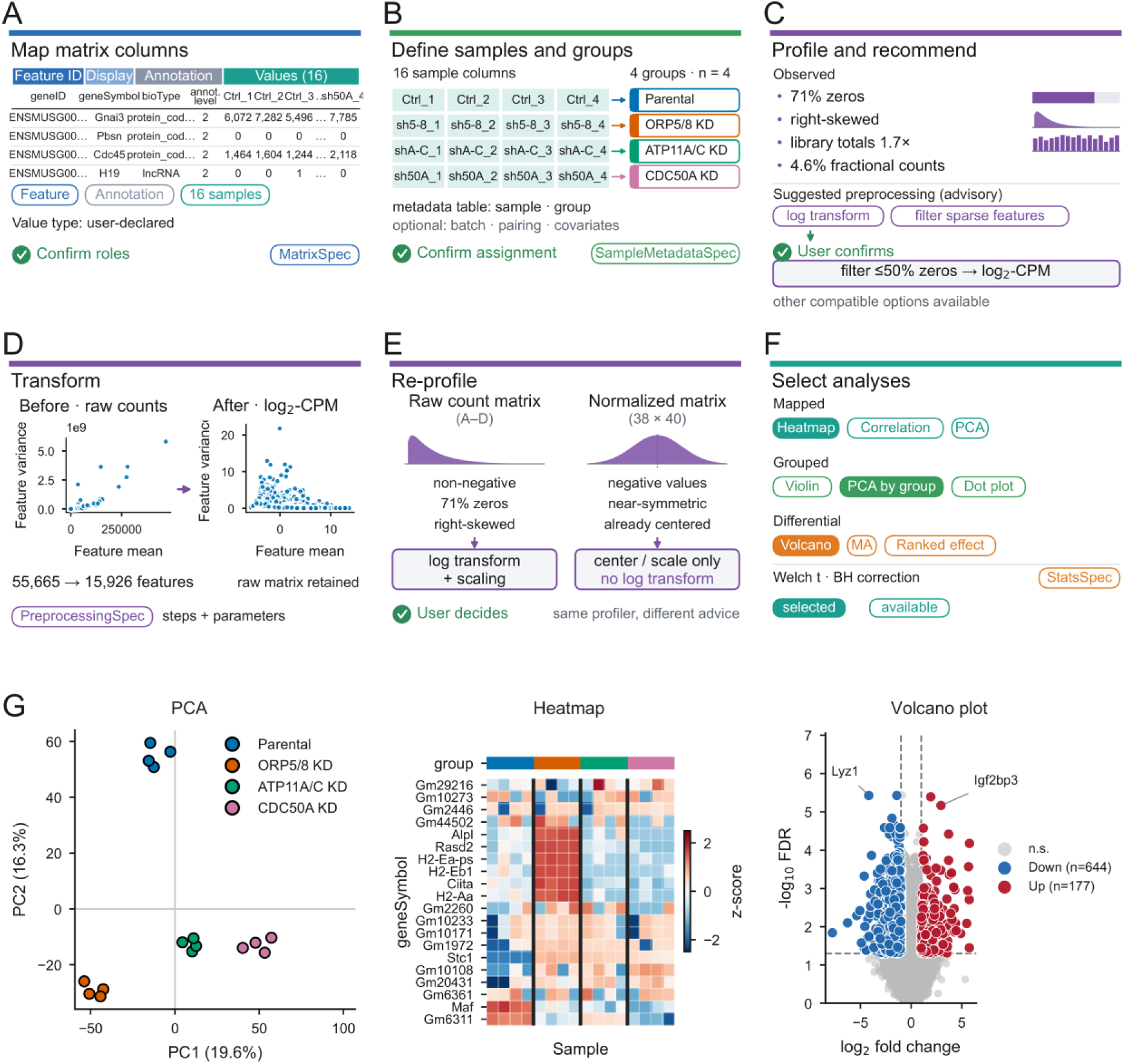
Guided matrix workflow from feature-by-sample input data to analysis-ready visualizations in MakeMyFigure. The workflow was demonstrated using the RSEM expected-count matrix of GEO series GSE299655 (55,665 genes × 16 samples; four groups of four replicates). (A) Matrix columns and data roles are identified and confirmed by the user. (B) Samples are linked to metadata and experimental groups. (C) Matrix properties are profiled, and preprocessing recommendations are provided. (D) User-selected preprocessing transforms the raw count matrix, with changes recorded in a reproducible workflow specification. (E) The profiler adapts recommendations to input data of different scales. (F) Statistical analyses are recommended and selected by the user. (G) Representative PCA, heatmap, and volcano plots show the analysis results.

Each stage produces a machine-readable JSON record (Fig. 1B). A MatrixSpec records which columns hold values and how the matrix is oriented, and a SampleMetadataSpec records the assignment of samples to groups and covariates. A PreprocessingSpec records the confirmed chain of transformations and filters that produced a derived matrix. A StatsSpec records the test, the comparisons, the correction, and the resulting values when a statistical test has been run. A PlotSpec records the plot type, column mapping, options, and aesthetics of one plot. A FigureSpec records a composite figure as a list of panels, each carrying its own PlotSpec, its StatsSpec, and the source table and worksheet it came from. A PlotSpec accompanies every specification export, and a StatsSpec is written whenever a test has been run; a composite carries its FigureSpec, and the PlotSpec of a plot derived from a matrix carries the identifiers of the matrix, metadata, and preprocessing records that produced it. Because the records are kept separate, the analytical and the visual decisions behind a panel can be inspected independently. Configuration intended for reuse is kept apart from these records. A Figure Preset stores the settings of a plot without its data, so that a laboratory or manuscript style can be applied to other figures of a compatible plot type, whereas PlotSpec and FigureSpec retain the exact configuration of one specific plot or composite (Fig. 3). A specification records the identity and a checksum of a source table rather than its contents, so regenerating a figure from a specification alone requires the data to remain available. A Figure Package removes this dependency: one portable file (.mmfpackage) that holds the specifications together with a frozen, lossless copy of the exact tables used, the original source file when available, the statistics and preprocessing records, imported images, and a manifest with a SHA-256 checksum of every file, so that the figure reopens on another computer without the original data.

**Figure 3.**
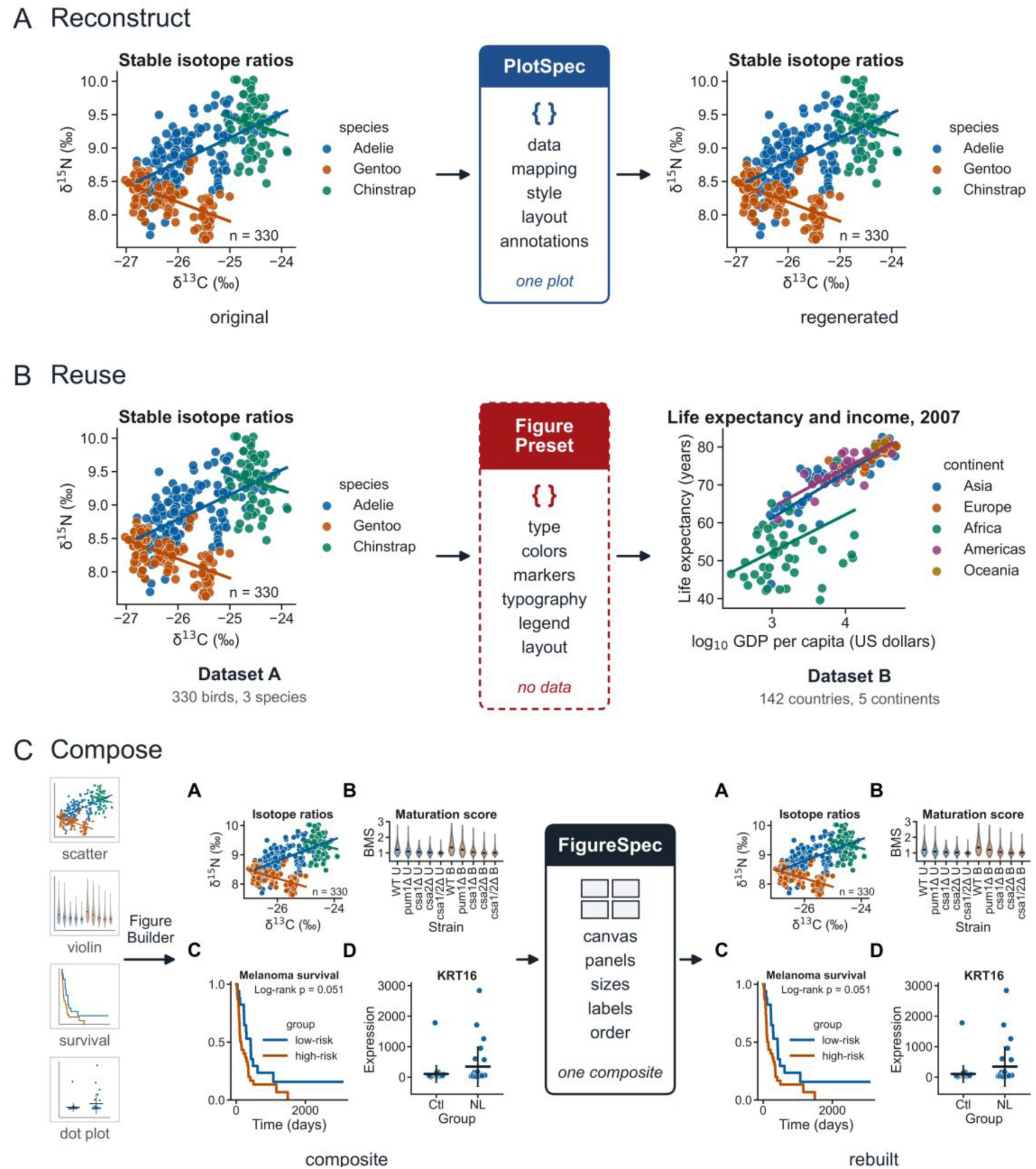
Figure specifications support reconstruction, reuse, and multi-panel assembly. (A) Reconstruct: a finished scatter plot saved as a PlotSpec is regenerated from that specification and its data table in a separate session, and the regenerated plot is identical to the original. (B) Reuse: a Figure Preset that carries only the appearance of a plot (colors, markers, typography, legend, and layout, but no data) is applied to an unrelated dataset, which keeps its own values, categories, and labels. (C) Compose: four panels are assembled with the Figure Builder into a composite, and the composite is rebuilt from its FigureSpec alone, with the same panel order, letters, sizes, and content.

MakeMyFigure therefore accepts two kinds of input: a table whose columns are already the variables of a plot, which can be drawn directly, and a feature-by-sample matrix, which must first be mapped, grouped and, where the data require it, transformed before it can be analyzed or plotted. The following section follows this second route on a deposited count matrix.

### A guided workflow takes a feature-by-sample matrix to analysis-ready plots

Many measurements arrive as a feature-by-sample matrix rather than as a table that is ready to plot, and such a matrix cannot be drawn until its columns have been told apart: which column identifies the feature, which columns annotate it, which columns hold the values of individual samples, and which samples belong to which biological group. MakeMyFigure handles this entry route in a dedicated matrix workflow (Fig. 2). For the gene-level count matrix deposited with GEO series GSE299655^23^ (55,665 genes × 16 samples with a gene identifier, a gene symbol, and two annotation columns), the workflow suggests the identifier and sample columns, with the option for the user to change them (Fig. 2A). The user confirmed the proposal, along with a declared value scale (never inferred), and it was recorded as a MatrixSpec. Samples were then assigned to their groups from a metadata table built from the GEO sample records, whose identifiers were matched to the value columns, and the assignment of replicates to each group was confirmed and recorded as a SampleMetadataSpec (Fig. 2B).

Before any transformation, the workflow profiles the confirmed values and shows what it observed: no negative or missing cells, zeros in 70.9% of cells, a strongly right-skewed distribution, per-sample totals of 24.0–41.5 million (a 1.7-fold range), and 4.6% non-integer cells, as expected for RSEM expected counts (Fig. 2C). From these properties a rule set suggests preprocessing chains that are compatible with the observed distribution, ranked and each accompanied by its assumptions and warnings; for this matrix the ranked suggestions were a log₂(x + 1) transform followed by row standardization, an arcsinh transform followed by standardization, and a sparse-feature filter followed by log₂(x + 1), with internal-standard and quantile normalization always offered. The suggestions are advisory and nothing is applied until the user confirms a chain. Here the user chose a count-aware chain from the method catalog, a filter that keeps genes with zeros in at most half of the samples followed by log₂ counts per million with a prior count of 0.5^24,25^. Instead of overwriting the original file, the program created a brand-new derived matrix containing the filtered genes, keeping the raw data completely untouched. Finally, it generated a PreprocessingSpec (Fig. 2D), a digital receipt that accurately recorded the recipe for the cleanup steps applied, ensuring the entire process is fully traceable and can be reproduced later without any guesswork. Profiling the derived matrix, or an independently deposited normalized matrix of transcription-factor activity scores (38 factors × 40 samples)^26^, produced the label “log-like or normalized” and a single suggestion to center or scale only, with no log transform offered (Fig. 2D,E).

Once the matrix was mapped and the sample groups are defined, MakeMyFigure suggests plots that are compatible for visualization based on available information (Fig. 2F). Matrix-level plots can be generated directly, whereas group-based plots require the sample groups and differential plots require a statistical comparison. The user can define the corresponding groups and statistical comparisons and utilize the differential plot visualization feature following the differential summary, which the user runs with a chosen test and correction (Fig. 2F). For example, Welch’s t-test with Benjamini–Hochberg correction^27,28^ identified 5,340 genes between CDC50A knock-down and parental cells at a false discovery rate below 0.05, including 821 genes with at least a twofold change. The resulting PCA, heatmap and volcano plot were generated through the same plotting workflow (Fig. 2G), with the data-processing and analysis steps retained with the exported figure. Together, these results show how MakeMyFigure can take a feature-by-sample matrix through mapping, preprocessing and statistical analysis to visualization, while leaving the choice of preprocessing and analysis with the user. The derived values, the Welch statistics and P values, the corrected P values and the PCA variances agreed with independent R calculations to within 10⁻¹³, or 10⁻⁸ for the corrected P values. Compared with count models run in R on the same contrasts (DESeq2, edgeR quasi-likelihood and limma-voom)^24,29,30^, the log₂ fold changes of the confirmed chain correlated at r ≥ 0.99 and the sets significant at 5% false discovery rate overlapped with Jaccard indices of 0.51–0.74, whereas the ranked log₂(x + 1) suggestion without library-size scaling reached only 0.01–0.46.

### Figure specifications support reconstruction, reuse, and multi-panel assembly

Once a plot has been generated and refined, MakeMyFigure records the settings needed to reconstruct it or reuse its appearance. A PlotSpec stores the data mapping, analysis and visual settings of one plot. To test reconstruction, we exported a scatter plot of stable-isotope ratios from 330 penguins^31^ and reopened its PlotSpec in a separate process (Fig. 3A). The regenerated plot reproduced the data points and fitted lines together with the colors, axes, labels, legend, annotations and dimensions, and all 47 PlotSpec fields were restored.

A Figure Preset serves a different purpose: it stores the appearance of a plot without its data. We saved a preset from the isotope scatter and applied it to an unrelated dataset of life expectancy and GDP per capita for 142 countries^32^ (Fig. 3B). The preset transferred the compatible colors, markers, typography, legend and layout, while the values, labels and categories came entirely from the new dataset. The log₁₀ transformation of income was recorded separately in a PreprocessingSpec. The same preset save-and-apply workflow was tested across all 38 registered plot types.

For multi-panel figures, the Figure Builder records the arrangement in a FigureSpec, including the canvas, panel positions and the specifications associated with each panel. We assembled four plots based on independent datasets^31,33–35^ and rebuilt the composite in a separate process (Fig. 3C). The rebuilt figure retained the panel number, order, labels, positions and dimensions. Because a FigureSpec records the identity of the source data rather than storing the data itself, reconstructing from the specification requires that the data remain available. For transfer between computers, MakeMyFigure therefore provides a Figure Package that stores the specifications together with a frozen copy of the required data. In testing, a packaged figure could be moved and reopened after the original data were removed, remained unchanged when the original file was subsequently modified, and failed its integrity check when the packaged data were altered.

### Outputs are consistent across input formats, computing environments, exports, data sizes, and repeated runs

MakeMyFigure is released for macOS, Windows, and Linux and accepts several input formats, so we next tested whether it produced consistent results across input formats and between Windows and Linux. Two datasets were loaded from the supported delimited-text and Excel formats and compared with their original tables (Fig. 4A). Delimited files returned identical values, whereas the small differences introduced through Excel were limited to numerical precision. We then ran the same data and specifications on Windows 11 and Linux under WSL2. Statistical results, plot coordinates, matrix values, and other numerical outputs were either identical or numerically equivalent between the two environments (Fig. 4B). Differences were confined to the placement of text elements that depend on font rendering or unseeded label positioning (Fig. 4C). Using the same font removed these differences except for repelled network labels. WSL2 was used for the Linux comparison; macOS was not tested.

**Figure 4.**
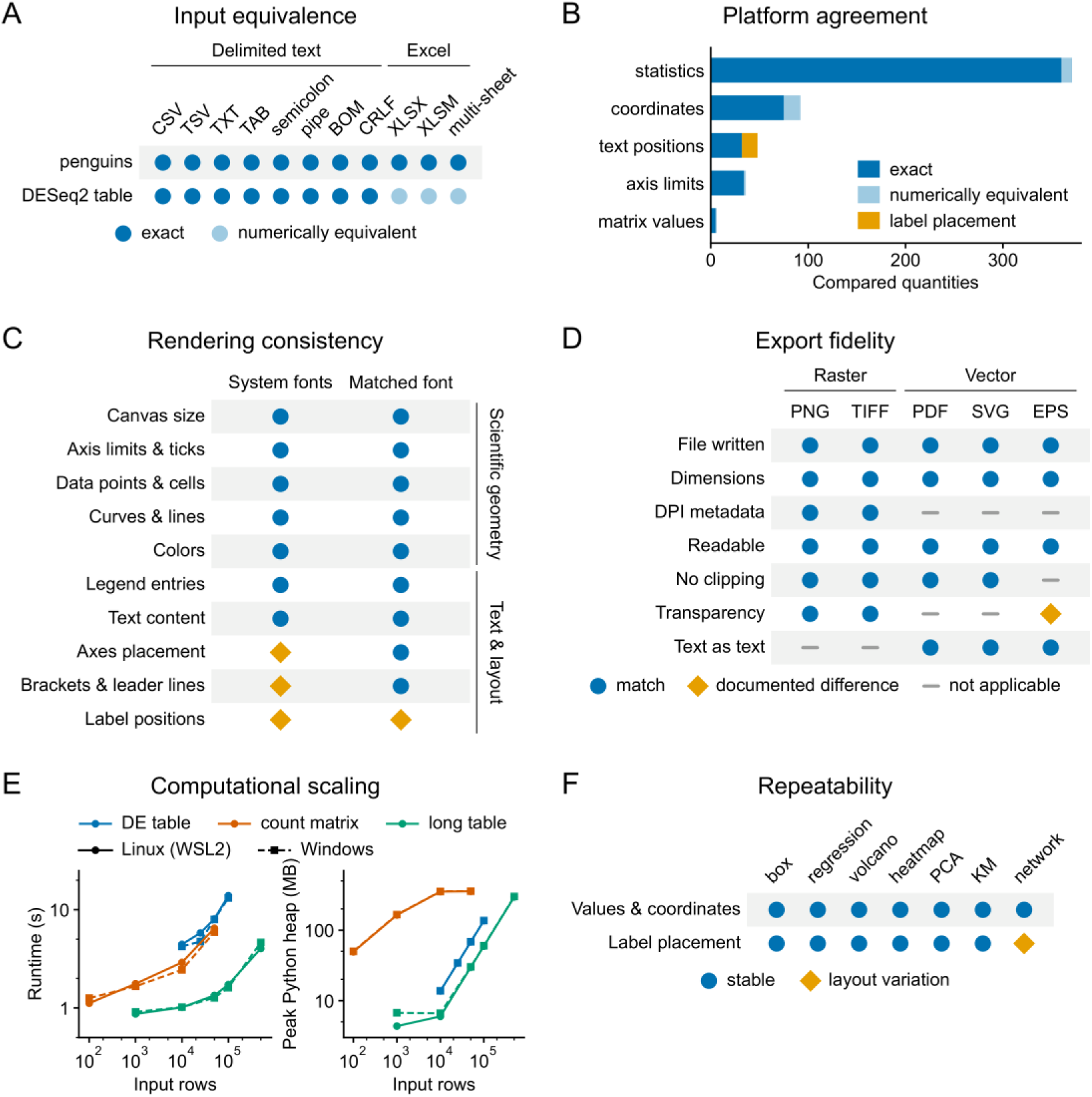
Consistency across input formats, computing environments, export formats, data sizes, and repeated runs. (A) The same two tables, supplied in eleven file encodings, load with identical values, or with numerically equivalent values for Excel files, which store fewer significant digits. (B) The comparison between Windows and Linux is exact or numerically equivalent; only the automatically placed text labels differ. (C) The scientific geometry of each plot is identical on both systems, and the text-dependent layout differs only when the systems use different fonts. (D) Every export-file property that applies to a format holds in all exported PNG, TIFF, PDF, SVG, and EPS files; EPS cannot store transparency. (E) Runtime and peak memory of the complete load, analyze, render, and export sequence for synthetic inputs of increasing size on both systems. (F) Twenty renders of each workload reproduce every value and coordinate, except for the shift in automatically placed labels on the network.

We also tested the five supported export formats (Fig. 4D). Raster and vector outputs retained the properties appropriate to each format, including dimensions, resolution for raster files and editable text for vector files. The documented exception was EPS transparency, which is not supported by the format. No clipping or unreadable exports were detected.

Finally, we measured runtime and memory use across differential-expression tables, count matrices, and long-format tables of increasing size (Fig. 4E). The largest tested input contained 500,000 rows, and runtime and memory increased with input size on both systems. Repeated rendering of the same specifications also preserved the numerical values and graphical elements across the tested workloads and plot types (Fig. 4F). The observed variation was limited to the placement of repelled text labels; the underlying data and graphical marks were unchanged. These benchmarks describe the tested workstation, environments, and dataset sizes rather than general performance across all hardware.

### MakeMyFigure recreates published figure panels from their source data

We next asked whether MakeMyFigure could reproduce figures from published biological studies using their deposited source data. We selected six open-access studies^26,36–40^ for which the values underlying a published panel were available in Source Data or supplementary tables (Fig. 5; Supplementary Fig. 1 and Supplementary Table 1). The examples span six journals and different biological questions and visualization types. All plots were generated directly from the deposited data; no values were digitized from published images.

**Figure 5.**
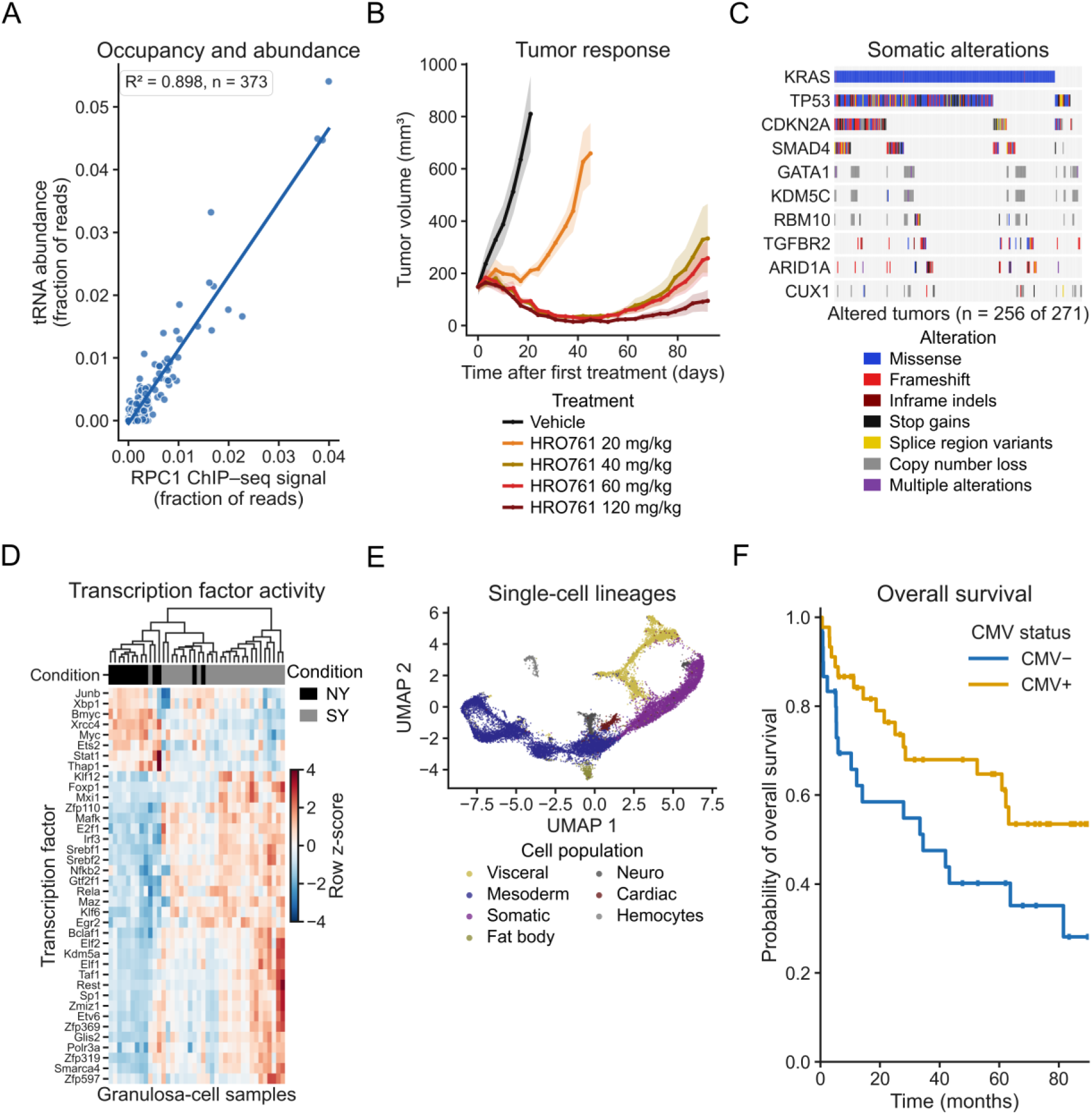
MakeMyFigure recreates figure panels from published source data. Each panel was regenerated from the source data deposited with an open-access article and assembled with the Figure Builder; Supplementary Table 1 lists the sources. (A) Scatter plot with regression: tRNA abundance against RNA polymerase III occupancy in human stem cells. (B) Time course: xenograft tumor volume in mice treated with vehicle or four doses of a WRN inhibitor (mean ± s.e.m., n = 5 mice per arm). (C) Oncoprint: somatic alterations in the ten most frequently altered genes across 271 pancreatic tumors. (D) Clustered heatmap: transcription-factor activity in granulosa cells from naturally ovulated and superovulated mice. (E) Embedding scatter: UMAP of Drosophila mesoderm and muscle nuclei colored by cell population. (F) Survival curve: overall survival of 75 melanoma patients receiving anti-PD-1 therapy, by cytomegalovirus (CMV) serostatus. Every plotted value comes from the deposited tables.

The examples include a regression between RNA polymerase III occupancy and tRNA abundance (Fig. 5A), tumor growth under increasing doses of a WRN inhibitor (Fig. 5B), an oncoprint of recurrent alterations in pancreatic cancer (Fig. 5C), and a clustered heatmap of transcription-factor activity in granulosa cells (Fig. 5D). The remaining examples show a UMAP of single-cell lineages in Drosophila (Fig. 5E) and Kaplan–Meier estimates^41^ of overall survival in patients with melanoma grouped by cytomegalovirus status (Fig. 5F). Each plot was generated from the corresponding published source table, and the six panels were assembled using the Figure Builder.

The recreated panels preserve the observations, transformations, and plot types used in the published figures, while allowing differences in visual styling and software-specific rendering. Side-by-side comparisons with the published panels are shown in Supplementary Fig. 1, and the source data and transformations used for each recreation are provided in Supplementary Table 1. Together, these examples show that the same MakeMyFigure workflow can be applied to published datasets with different structures and visualization requirements.

### Independent implementations validate statistical calculations, and published analyses are reproduced from source data

Next, we asked whether the statistics produced by MakeMyFigure agreed with those calculated independently in R (Fig. 6A). Both implementations used the same input data and the same statistical procedure. Nine representative methods are shown in the main figure, spanning two-group tests, correlation, analysis of variance, regression and survival analysis, with the complete benchmark provided in Supplementary Fig. 2. Across 22 benchmark datasets, all 143 test statistics, P values and effect-size or slope estimates agreed with R within the prespecified tolerance (Fig. 6B). Across the complete benchmark of 2,625 quantities, none failed the comparison; the remaining differences reflected documented numerical or methodological conventions rather than disagreement in the underlying calculations.

**Figure 6.**
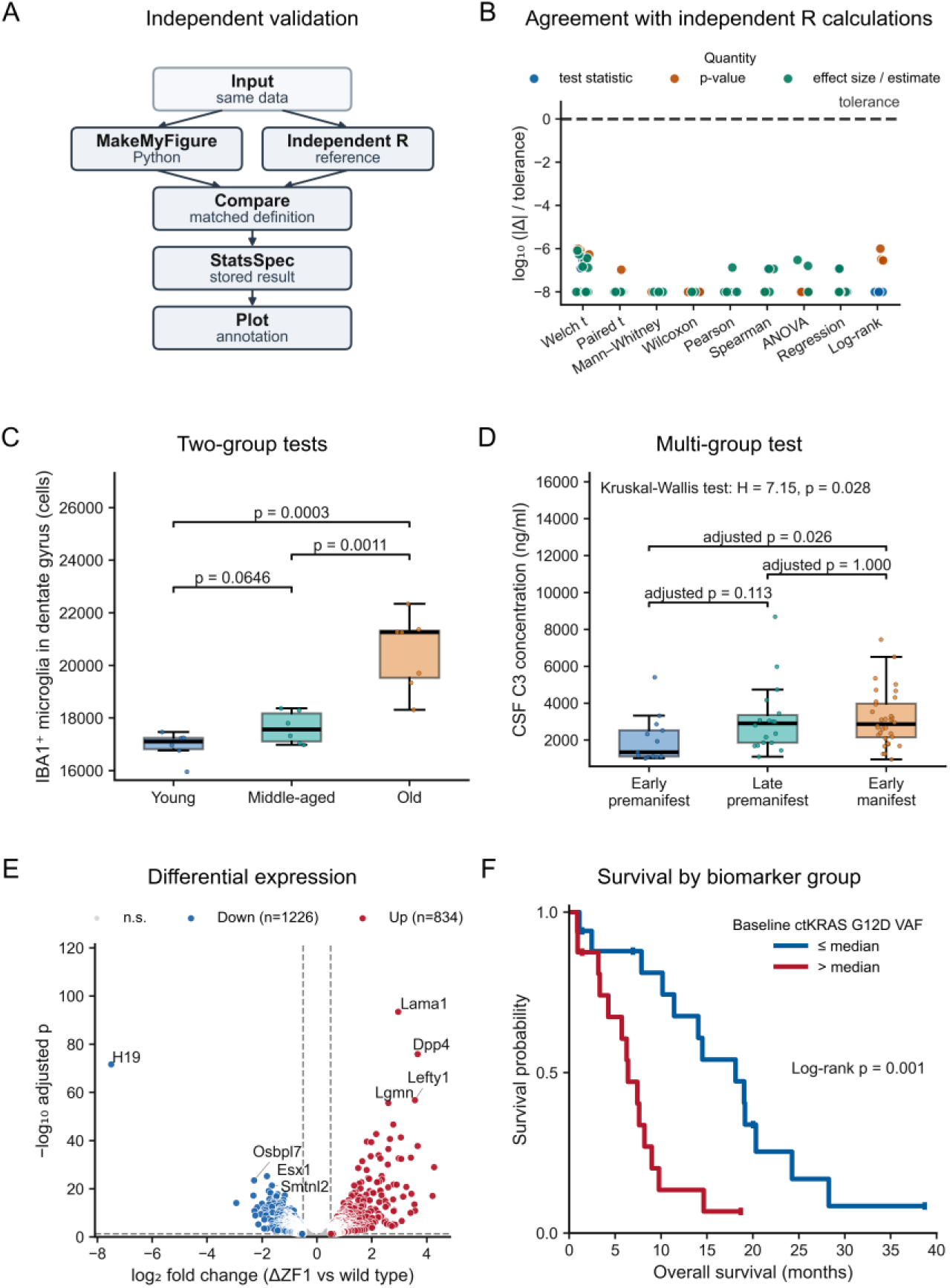
Independent implementations validate statistical calculations and reproducibility. (A) The same tests were run by MakeMyFigure and independently written R code, and the results were compared; figure annotations come only from the stored StatsSpec. (B) Agreement with R for nine representative procedures: each point is one compared statistic, P value, or effect estimate, shown relative to its tolerance. (C) Two-group tests were conducted on the published data to replicate the statistical comparison; the P values shown are equal to the published Welch t-test P values. (D) Multi-group tests were conducted on published data to demonstrate the reproducibility of the statistical analysis results from the authors’ Kruskal–Wallis test and Dunn’s post hoc test; the adjusted P values were found to equal the published values. (E) A published DESeq2 result table was plotted as a volcano plot using the article’s thresholds, with exact counts of up-and down-regulated genes matching those reported. (F) Kaplan–Meier curves for 33 pancreatic cancer patients, split at the median circulating KRAS G12D fraction, were drawn in MakeMyFigure; the log-rank P value it computed matches the published value. Boxes in (C) and (D) show the median and interquartile range, whiskers extend to 1.5 × IQR, and points are individual samples.

We then applied the same framework to statistical analyses reported in published studies. For IBA1⁺ microglia counts in young, middle-aged and old mice^42^, pairwise Welch *t*-tests gave P = 0.0646, 0.0011 and 0.0003, matching the published values and independent R calculations (Fig. 6C). For cerebrospinal-fluid complement C3 across three stages of Huntington’s disease,^43^ a Kruskal–Wallis test followed by Dunn’s test with Bonferroni correction gave adjusted P values of 0.113, 0.026, and 1.000, again matching the published values at their reported precision and the independent R calculations (Fig. 6D).

MakeMyFigure can also visualize differential-expression results generated by specialized RNA-seq pipelines without recalculating them. A published DESeq2 table comparing CTCF ΔZF1 with wild-type neural progenitor cells^44^ was mapped to gene, log₂ fold change, and adjusted P value and plotted using the thresholds reported in the source study (Fig. 6E). This identified 834 up-regulated and 1,226 down-regulated genes, matching the published counts. This panel, therefore, demonstrates visualization of imported differential-expression results rather than agreement between MakeMyFigure and DESeq2.

Finally, we reproduced a published survival analysis of 33 patients with metastatic pancreatic ductal adenocarcinoma grouped by baseline circulating KRAS G12D variant-allele fraction^45^. The log-rank test (P = 0.0010) and Cox hazard ratio of 4.18 (95% CI, 1.68–10.39) agreed with independent R calculations and the published results (Fig. 6F). Together, these analyses show that statistical results generated within MakeMyFigure can be independently reproduced, while results produced by external specialist methods can be imported and visualized without being recalculated (Supplementary Table 2).

## Discussion

Figure preparation is often the final step of a quantitative study, yet the tools used for statistical analysis, visualization, and figure assembly have largely developed as separate applications. MakeMyFigure was developed to centralize these operations and, more specifically, to record each operation associated with the panel it produced. The present work shows that a single local application can connect these steps for tabular data. It took a deposited count matrix from column mapping to analysis-ready plots through a profiled, confirmed and recorded preprocessing chain (Fig. 2). A plot, a reusable style, and a four-panel composite were regenerated from their recorded specifications in separate processes (Fig. 3). Numerical output remained consistent across file encodings and two operating environments, while rendering was evaluated across five export formats and twenty repeated renders, with the observed differences confined to text placement (Fig. 4). MakeMyFigure also recreated six published panels representing six visualization classes from their deposited source data (Fig. 5), and its statistical procedures agreed with independent R implementations within prespecified tolerances (Fig. 6).

Of the tools compared in Table 1, GraphPad Prism^8^, OriginPro^9^, JMP^10^, SigmaPlot^11^, and DataGraph^12^ combine analysis with publication graphics in mature desktop applications, with capabilities such as Prism’s assumption checks, JMP’s persistent row states, and SigmaPlot’s curve-fitting library that MakeMyFigure does not reproduce. jamovi^13^ and JASP^14^ provide point-and-click statistical analysis, including Bayesian methods in JASP, whereas Perseus^15^, iDEP^16^, and ExpressAnalyst^17^ provide workflows designed around omics matrices. MakeMyFigure instead performs standard statistical analyses within the figure workflow and can import results from specialist methods, including count-based differential-expression pipelines, rather than attempting to replace them.

Other tools address different parts of the workflow. Galaxy^18^ and KNIME^21^ preserve computational analyses as re-runnable workflows, SRplot^19^ and Hiplot^20^ provide broad collections of scientific visualizations, and BioRender^22^ combines figure preparation with data import and statistical functions. Code-based environments such as ggplot2 and^46^, matplotlib^47^, together with assembly tools such as plotgardener^48^, provide extensive control and reproducibility through scripts, while grammars such as Vega-Lite^49^ demonstrate how visualizations can be represented as machine-readable specifications. The distinction in MakeMyFigure is the integration of these steps in a code-free workflow in which the source data, confirmed preprocessing, statistical results, plot configuration, and multi-panel assembly remain linked through records that can be reopened within the same application. It therefore complements rather than replaces specialist statistical, omics, and visualization tools.

A central part of the design is the record retained with each step of the figure workflow. A PlotSpec records the data mapping and settings needed to regenerate a plot from its source table, while a StatsSpec records the test, comparison, and correction associated with statistics shown on the figure. A PreprocessingSpec records transformations without modifying the original data, a Figure Preset stores reusable visual settings without the data, and a FigureSpec records the organization and source of panels in a composite (Fig. 3). Keeping these records separate allows the analytical and visual decisions behind a figure to be inspected independently, and their JSON format makes them readable outside the application.

A specification alone does not contain the underlying data and therefore requires the recorded source table to regenerate a figure. The desktop application checks the source against its recorded checksum and warns if it has changed. For transfer between computers, a Figure Package stores the specifications together with a frozen copy of the required data and verifies their integrity with checksums. Thus, the specifications provide an inspectable record of how a figure was generated, whereas the Figure Package provides the data and records needed to reopen it independently of the original files. This is critical for the reproducibility of work.

This work has several limitations, some inherent to the design and others defined by what was tested. The recommendations are rule-based and advisory: they suggest plots, transformations and statistical tests from the detected structure of the data, but the researcher remains responsible for selecting an appropriate analysis and evaluating its assumptions. The same applies to preprocessing. For example, the initial recommendations for the count matrix in Fig. 2 did not include the library-size scaling required for that dataset, and the count-aware preprocessing chain was selected by the user.

Rendering can vary slightly across systems because font metrics affect axis and label placement, and three plot types use an unseeded label-repulsion procedure that can change label positions between renders without changing the underlying data marks (Fig. 4C,F). Cross-platform testing was performed on Windows and Linux under WSL2 on a single workstation; native Linux and macOS were not tested, and the performance measurements apply only to the hardware and dataset sizes examined here.

The Figure Builder uses a grid-based layout and does not currently support free positioning or layering. A FigureSpec records a composite but requires its associated data to reconstruct it, whereas a Figure Package carries the required data and can be reopened through the application. The published-panel examples are scientific rather than exact reproductions because the original graphics software and, for the heatmap, the original clustering implementation were not rerun, and some statistical decorations from the published panels were not reproduced. Finally, the Publication style provides a general manuscript-oriented format rather than guaranteeing compliance with the requirements of a particular journal.

Future development will follow the principle that shaped this version: new capabilities should be accompanied by a record that can be inspected and validated. Because MakeMyFigure runs locally, data can remain within the research environment in which they are generated. By retaining the data and records associated with a figure, the application allows analyses and figures to be revisited, corrected, and reconstructed after the original session has ended.

## Conclusion

MakeMyFigure brings data ingestion, confirmed transformation, statistical testing, plotting, annotation, and multi-panel assembly into a single code-free application, while retaining a machine-readable record of each step. These records and the associated data can be stored together in a portable Figure Package. Using public data, MakeMyFigure took a deposited count matrix through a confirmed preprocessing workflow to analysis-ready plots, recreated published panels from their source data, and produced statistical results that agreed with independent R calculations within prespecified tolerances. Plots, styles, and multi-panel figures were also regenerated from their recorded specifications, and outputs remained consistent across the tested input formats, computing environments, export formats, and repeated renders within documented limits. The software, documentation, and benchmark materials are openly available so that these results can be reproduced and extended.

## Methods

### Software implementation

MakeMyFigure is written in Python (3.9 or later) as a frontend-agnostic core package, make_my_figure_core, and two graphical interfaces that call the same functions: a desktop application (PySide6) and a local browser application (Streamlit). The core depends on pandas^50^, NumPy^51^, SciPy^52^, statsmodels^53^, matplotlib^47^, openpyxl (https://openpyxl.readthedocs.io), jsonschema (https://python-jsonschema.readthedocs.io), NetworkX^54^, adjustText^55^, and Pillow (https://python-pillow.github.io); no R installation, network service, or account is required, and no data leave the machine. Data flow through one path in both interfaces: a loader produces a table with column classifications; a validated PlotSpec (JSON Schema draft 2020-12, https://json-schema.org/draft/2020-12) selects one of 38 registered renderers; the renderer receives the table, the specification, and a resolved style profile and returns a matplotlib figure with metadata and warnings; the export layer writes the requested formats, a sidecar record on specification exports, and, on request, a figure package. Every renderer draws from the same style profile (45 typographic, geometric, and color tokens), so all plot types share a single visual language, named Publication; it is a general, manuscript-ready style and does not claim compliance with any journal. The version described here is release 1.1.0 (release tag v1.1.0, 6 September 2026); the figure package (Results, Methods) is implemented on the repository branch feature/portable-figure-package-v1.1.1 (commit 193f637), and the technical tests of Fig. 4 were run on release 1.1.0; release installers for Windows, macOS, and Linux are built with PyInstaller and require no Python installation.

### Data intake and column mapping

Delimited text (comma, tab, semicolon, or pipe; .csv, .tsv, .txt, .tab, with or without a byte-order mark and with either line ending) is read with pandas after the delimiter is sniffed from the first lines; numeric conversion uses the round-trip float converter so that a decimal string is parsed to the nearest double. Excel workbooks (.xlsx, .xlsm; legacy .xls through an optional reader) are read with openpyxl; every worksheet is listed with a heuristic classification (differential results, matrix, metadata, enrichment, documentation, empty, generic), any sheet can be chosen, header rows can be selected or combined, and merged header cells are recovered from the workbook’s merge map. Columns are classified as numeric or categorical after coercion; identifier-like columns are kept as text. A table can also be reshaped in the application without a metadata file: a wide features-by-samples matrix can be melted to a long table with an in-app sample-to-group assignment, and a grouping column can be derived from an existing column. Column roles required by a plot type (for example x, y, group, time, event, P value) are proposed from the classification and from case-insensitive alias lists for differential-expression headers (edgeR/limma and DESeq2 conventions). They are always confirmed or changed by the user before anything is plotted, and no column is ever substituted silently.

### Data profiling and recommendations

A rule-based engine profiles the table (column kinds, identifier columns, binary event columns, matrix blocks) and assigns it to one of 15 input schemas (for example precomputed differential results, survival, dose–response, network edge list, mutation matrix, enrichment, expression-like matrix, matrix with metadata, paired, generic long table). For the detected schema the engine lists candidate plot types with a draft column mapping and a one-line reason, compatible statistical tests for the chosen plot and mapping, and compatible transformations, each with a warning where an assumption is not met. The recommendations are advisory: nothing is applied until the user selects it, and the selection is recorded in the PlotSpec. No machine learning is used.

### Matrix workflow, preprocessing, and transformation records

Feature-by-sample matrices are handled in a dedicated workflow with steps including mapping value columns (MatrixSpec), assigning samples to groups (SampleMetadataSpec), preprocessing, validation, and plot generation. Preprocessing steps are log_2_, natural log, and log_10_ with a pseudocount, arcsinh, square root, winsorization, imputation (feature median, constant), feature filtering by missingness or intensity, scaling and normalization (total-sum scaling and counts per million, median scaling, upper-quartile scaling, quantile normalization (the Bolstad et al. algorithm as implemented by limma’s normalizeQuantiles)^30,56^, trimmed mean of M-values (TMM)^57^, the voom log_2_-CPM transform of limma without precision weights^25,30^, row or column z-scores, robust scaling, centering, and normalization by internal-standard features or a reference sample), and derived views (top-variance features, group means, sample and feature correlation, distance matrices). Quality-control summaries report per-sample totals, missingness and distribution shape and indicate whether the matrix has count-like or intensity-like characteristics.

Every preprocessing step is shown with its parameters and is applied only after user confirmation. The original matrix is retained and the transformation is applied to a derived copy. The ordered preprocessing chain, including method names, parameters, input and output dimensions and matrix identifiers, is recorded in a PreprocessingSpec. PlotSpecs generated from the derived matrix reference this record, and a Figure Package contains the PreprocessingSpec together with the source and derived matrices. For Fig. 3B the Gapminder 2007 table (142 countries)^32^ was transformed by log_10_ of the gdpPercap column through this workflow; the SHA-256 of the raw file before and after the run was compared with the value recorded before any run, and the derived values were recomputed independently with numpy.log10.

Profiling of the confirmed values computes the numbers of features, samples and missing cells, the fractions of zero and negative cells, whether every cell is integral, the range, mean and median, per-feature variances, the skewness of all values and of each sample (SciPy, biased estimator), per-sample totals and medians, and outlier samples or features whose total or variance lies more than 3.5 median absolute deviations from the median. From these a label is attached (count-like if all cells are integral, non-negative, right-skewed and the maximum is at least 100; intensity-like if non-negative and skewed; log-like or normalized if negatives lie within ±40 or if |skewness| < 1 and the maximum is below 40) and preprocessing chains are suggested by fixed rules: center or scale only for log-like matrices; total-sum or median scaling followed by log₂ when per-sample totals differ more than threefold; log₂(x + 1) or arcsinh followed by row standardization when skewness exceeds 1.5; a sparse-feature filter followed by log₂ when more than 30% of cells are zero; standardization for visualization only when negatives are present; and, always, internal-standard and quantile normalization with their caveats.

These labels and recommendations are advisory and are shown with the rule that produced them; the graphical wizards apply a suggested chain as a whole after confirmation, and any ordered chain of the 27 registered methods can be applied through the Python interface. Quality-control plots, including per-sample totals, medians and distributions, zero and missing fractions, sample correlations, PCA and mean–variance trends, can be generated before and after preprocessing.

The feature-level differential summary compares confirmed groups on the derived matrix with Welch’s or Student’s t-test, the Mann–Whitney U test, paired t or Wilcoxon signed-rank tests, one-way ANOVA or the Kruskal–Wallis test, with Benjamini–Hochberg, Bonferroni or Holm correction. It reports group means and medians, log₂ fold change, and an effect size appropriate to the selected test. Log₂ fold change is calculated as the mean difference for data on a confirmed log scale, as the log₂ ratio of pseudocount-adjusted means for data on a linear scale, and as a mean difference when the scale has not been confirmed. Effect sizes are Cohen’s d with a t-based 95% confidence interval or rank-biserial correlation, as appropriate. The output also records the preprocessing chain used for the comparison. This procedure is not a count model; an optional negative-binomial analysis using PyDESeq2^58^ is available as a separately installed extra (count-de) and was not used here.

Plot recommendations become available as the information required for them is defined: matrix plots after matrix mapping, group comparisons after groups are assigned, and volcano, MA and ranked-effect plots after a differential summary is generated. Selected recommendations are constructed through named transformations, including wide-to-long reshaping, row standardization, top-variance selection, sample or feature correlation and PCA of centered values, and are opened in the plot editor with references to the associated matrix, metadata, preprocessing and statistical records.

For Fig. 2 the GEO GSE299655 supplementary file^23^ (RSEM expected counts, 55,665 genes × 16 samples; SHA-256 identical to the deposit) was mapped as proposed with the value scale declared as raw counts, and the four groups of four replicates were taken from the GEO sample records. The confirmed chain was a filter keeping genes with zeros in at most 50% of samples and non-zero variance (15,926 genes retained) followed by log₂-CPM with a prior count of 0.5; the differential summary used Welch’s t-test with Benjamini–Hochberg correction for each knock-down against the parental line. The transcription-factor activity matrix of Daugelaite et al.^26^ (Source Data Fig. 3d; 38 × 40; values as deposited) was mapped with the value scale declared as normalized and profiled without applying any step.

The preprocessing chain, statistical tests and the PCA were recomputed independently in R 4.3.3 (edgeR cpm, t.test, p.adjust, prcomp), and the differential summary was compared with DESeq2, edgeR quasi-likelihood and limma-voom results for the same contrasts from the R validation by the Pearson correlation of log₂ fold changes, the direction agreement of genes significant in both, the overlap of the 100 smallest P values and the Jaccard index of the sets significant at 5% false discovery rate. Panels A–F were drawn with matplotlib primitives from the workflow’s JSON records; panels D and G are MakeMyFigure renders opened from the workflow (heatmap row labels switched to gene symbols and two volcano labels selected in the plot editor), and the figure was composed with PyMuPDF as for Fig. 3.

### Statistical analysis and multiple-testing correction

Eighteen statistical procedures are registered in a single engine used by both interfaces. The user selects the test and comparison family for a plot. Available methods include Student’s, Welch’s and paired t-tests, the Mann–Whitney U test^59^ and the Wilcoxon signed-rank test^60^; one-way, two-way, and repeated-measures analysis of variance; Kruskal–Wallis test^61^, with Dunn’s test^62^; χ^2^;^63^ and Fisher’s exact tests^64^; log-rank test^65^ and Cox proportional-hazards analyses^66^; Pearson and Spearman^67^ correlation; and linear and generalized linear regression.

Tests are two-sided, with Welch’s t-test^27^ being the default two-group test. Paired tests match observations by subject identifier, and missing observations are omitted within the relevant comparison. P values within a comparison family are adjusted together using the Benjamini– Hochberg^28^, Holm^68^ or Bonferroni^69^ procedure.

Each result is stored in a StatsSpec containing the test, comparisons, correction, statistic, P and adjusted P values, effect size and sample sizes. Statistical annotations are drawn from these stored results rather than recalculated by the renderer. Core implementations use SciPy and statsmodels, with selected procedures implemented within MakeMyFigure; detailed implementation conventions are provided in the Supplementary Methods. Precomputed results, including differential-expression tables, are plotted from their mapped source columns and are not recomputed.

### Differential-result handling

MakeMyFigure computes differential expression natively through the feature-level differential summary described above, and a negative-binomial count model is available as an optional extra; precomputed results produced by dedicated count-model tools are handled separately and are never recomputed. A table produced elsewhere, for example by DESeq2^29^, edgeR^24,70^ or limma^30^, is mapped to roles (log₂ fold change, P, adjusted P, mean abundance, identifier) and drawn as a volcano or MA plot with user-specified thresholds; the renderer counts features that pass the thresholds with inclusive inequalities and reports the counts, and the published values are never modified. For Fig. 6E, table S4 of Lucero et al. 2025^44^ (Europe PMC supplementary files, SHA-256 recorded; DESeq2 results for NPC ΔZF1 versus NPC wild type; 55,291 genes with the columns gene, baseMean, log2FoldChange, lfcSE, pvalue and padj) was mapped to gene, log₂ fold change and adjusted P and drawn with the thresholds of the article’s Fig. 4A legend (adjusted P ≤ 0.05, |log₂ fold change| ≥ 0.5). The 40,502 genes without an adjusted P value, which DESeq2 excludes by independent filtering, were reported by the renderer and not drawn; no gene lay on a threshold; the four most significant genes in each direction were labeled by adjusted P with one label per symbol. The counts of 834 up-and 1,226 down-regulated genes were re-evaluated independently with pandas and compared with the counts stated in the article’s Results. For matrices with no prior analysis the application performs its own differential analysis, in the benchmark a per-feature screen (Welch’s *t* on log₂ counts per million with a prior count of 0.5 and Benjamini–Hochberg correction^28^), labeled in the interface as a screen to distinguish it from a count model; its agreement with the same procedure written in R is part of the benchmark above (100 comparisons), and its concordance with limma-voom, edgeR and DESeq2 on a public count matrix is described in the Supplementary Methods (Supplementary Fig. 3).

### Plot rendering, annotation, and export

Renderers never modify the input table (they operate on a copy) and take fonts, line widths, marker sizes, palette, and legend placement from the style tokens; the user can override any token or layout option (column width, aspect, margins, legend location, axis scales) in the PlotSpec. Point-level annotation supports text, arrows, callouts, boxes, regions, brackets, and reference lines in data, axes, or figure coordinates, recorded in the PlotSpec. Automatic label placement in the volcano, lollipop, and network renderers uses adjustText, whose repulsion search is not seeded; these are the three plot types whose text positions may differ between renders. Figures are exported as PNG and TIFF (at a chosen resolution, 300 dpi by default) and as PDF, SVG, and EPS with text kept as text (TrueType type 42 embedding; SVG text elements). Specification exports (the PlotSpec export, the export bundle, and the browser downloads) write a sidecar file containing the PlotSpec, the render metadata (software versions, columns used, counts, warnings), and, when statistics were run, the StatsSpec; the single-format buttons write the figure file alone. The PlotSpec records a SHA-256 content digest of the table it was drawn from, and the desktop application warns when a PlotSpec is opened against a table with a different digest.

### PlotSpec, Figure Presets, FigureSpec, and Figure Packages

A PlotSpec records one plot: its plot type, source table and, for multi-sheet workbooks, workbook and worksheet; column mapping; plot options and thresholds; style and layout; annotations; statistical configuration; and export settings. A Figure Preset (format version 1, .mmfpreset.json) stores settings intended for reuse without storing the underlying data. In style mode it retains typography, palette, geometry, legend and colorbar placement, export size and plot-specific visual options. Full mode additionally retains column roles, thresholds, axis labels and the statistical test. Presets contain no table values, labels or annotations, and the validator rejects presets containing data. When a preset is applied, compatible settings are transferred to the new PlotSpec and any column roles that cannot be resolved in the new table are reported for user mapping. The same implementation is used by both interfaces.

A FigureSpec (.figure_spec.json) records a multi-panel composite, including its canvas dimensions, grid, spacing, panel-letter style and embedding resolution. For each panel it records the label, size, PlotSpec, StatsSpec and source table or worksheet; imported images are recorded with their SHA-256 checksum, dimensions and fit mode. The Figure Builder arranges panels on the recorded grid while preserving their aspect ratios. Generated panel content is embedded as raster at the specified resolution, while panel letters and titles remain vector elements. Composites can be exported as PNG, PDF and SVG. A Layout preset stores the grid, panel sizes, spacing and letter style without panel content.

A Figure Package (format version 1, .mmfpackage) is a portable ZIP container written and read by the core library and supported by both interfaces. It contains the PlotSpec of a single plot or the FigureSpec of a composite together with the PlotSpec and StatsSpec associated with each panel. Required tables are stored in a lossless representation that preserves data types, row and column order, missing values and floating-point values; when accessible, the original CSV, TSV or Excel file is retained as well. For matrix-derived plots, the package also contains the source and derived matrices together with the MatrixSpec, SampleMetadataSpec and PreprocessingSpec. Imported panel images, PNG, SVG and PDF previews, software-environment information and a manifest listing every file with its size and SHA-256 checksum are also included. Identical tables are stored once, and absolute source paths are not retained.

When a Figure Package is opened, the container is checked for unsafe entries, the manifest is validated against its JSON Schema, and every listed file must be present and match its recorded checksum before rendering begins. Packages with altered contents or an unsupported newer format version are rejected. The figure is then rendered from the frozen data. Stored statistics are checked against the StatsSpec, and recorded preprocessing can be replayed from the frozen source matrix and compared with the stored derived matrix without replacing the frozen values. Because a Figure Package contains data, both interfaces state this before saving. A standalone PlotSpec instead records the SHA-256 digest of its source table; when opened in the desktop application, the user is warned if the selected table no longer matches that digest.

### Reconstruction, reuse, and composition round trips

For Fig. 3, all stages ran as separate Python processes launched by one driver script so that no regenerated object shares memory with its original. Reconstruction: a finalized scatter of stable-isotope ratios by species (Gorman et al. 2014, 330 birds; palmerpenguins release, CC0)^31^ was exported with its PlotSpec sidecar; a new process loaded the PlotSpec through the desktop application’s loader, resolved the table by its recorded name, rendered, and exported again. Structural comparison recorded the coordinates of every point collection and fitted line, face colors, axis limits, tick labels, axis labels and title, legend entries and location, annotation text and position, font sizes, spine visibility, and figure size; the SVG drawing commands after removal of metadata, dates, and volatile element identifiers; and the PNG pixels; all were compared for exact equality, and PlotSpec fields were compared leaf by leaf (47 fields). Reuse: a style preset and a full preset were extracted from the finalized PlotSpec and stored; a new process loaded the Gapminder 2007 table, derived log_10_ of income per person through the matrix workflow, built a default PlotSpec naming the new table and columns, applied the stored style preset, and rendered; the full preset was applied as a negative control and its report of unresolved roles recorded. The same extract–apply–render sequence was run for every registered plot type from its bundled example (38 of 38 passed). A separate audit compared 39 preset properties at the source, serialized, and applied levels (Source Data). Composition: four generated panels were assembled by the Figure Builder (two columns, 180 mm, per-panel widths, letters A–D, 600-dpi embedding) and the FigureSpec written. The panels were the isotope scatter, basidial maturation scores from Liu et al. 2018^33^, the GEO GSE51547 survival table from Guo et al. 2019 with a log-rank test^34^, and KRT16 expression from Kume et al. 2024^35^. A new process rebuilt the composite from the FigureSpec alone, re-attaching each table by its recorded name. The number of axes, canvas size, per-panel cell position and size, letters, panel order, and SHA-256 of each embedded panel raster were compared, together with eight builder geometry checks (resizing, aspect preservation, equal widths, reordering and re-lettering, width ratios, imported-asset record, export formats, letters drawn). The transformation record was checked as described above. Figure packages were tested in the automated test suite (61 tests): for scatter, statistical box and bar, volcano, clustered heatmap, PCA, Kaplan–Meier with log-rank, forest and network plots, a package was written from a rendered figure and reopened, and a structural signature of the reopened figure (data coordinates, axis limits, ticks, labels, colors, legend, size and, where run, every statistic) was compared with the original; in a separate-process test a statistical box plot was packaged, the package moved to a new directory, the original data directory deleted, and the package reopened in a clean Python process from the package alone; editing the original file after packaging left the reopened figure unchanged; altering one value inside a package, replacing a checksum, adding an unlisted file, removing a file, corrupting the manifest, declaring a newer format version, and container entries with path traversal, links or extreme compression were each refused. All 38 plot types were packaged and reopened, and a matrix-derived heatmap package replayed its recorded preprocessing chain to the frozen derived matrix exactly.

### Input-format equivalence, cross-platform, export, performance, and repeatability tests

For Fig. 4, two real tables, the penguin morphometrics (344 × 8)^31^ and the complete DESeq2 result table of Osipovich et al. 2023 (17,028 × 8; S2 Table)^71^, were re-encoded with pandas and openpyxl as comma-, tab-, semicolon-, and pipe-delimited text (.csv, .tsv, .txt, .tab), CSV with a UTF-8 byte-order mark, CSV with CRLF line endings, single-sheet XLSX, XLSM, and a five-sheet XLSX with the data sheet in second position, and loaded through the same loader; shape and column names, every numeric cell (bit identity, maximum absolute and relative difference, missing-value pattern), categorical values and their order, the column classification, the column mapping, and the coordinates of the plot rendered from each file were compared with the original CSV (tolerance 10^−12^ relative; legacy .xls not tested). The cross-platform comparison ran seven workloads (box plot with Welch tests and Benjamini–Hochberg correction; per-group regression; volcano; clustered heatmap and PCA of GEO GSE299655 after CPM filtering, log₂ CPM, top-variance selection, and row z-scores; Kaplan–Meier with log-rank; seeded network layout) from identical data and specifications on Windows 11 (build 26200) and on Ubuntu 22.04 under WSL2 on the same Intel Xeon Silver 4208 processor, with identical Python library versions (numpy 2.1.2, pandas 2.2.3, SciPy 1.17.1, statsmodels 0.14.6, matplotlib 3.10.8, Agg backend), under two font arms (each system’s own fonts, Arial versus DejaVu Sans; and DejaVu Sans on both). For each render the suite recorded, without screenshots, the figure size, axis limits, scales, ticks, and labels, legend entries, every line’s data, every collection’s offsets, sizes, and colors, every patch’s vertices, every text’s string and position, and every image array, stored them as arrays and compared them element-wise, allowing a per-component sign flip for PCA; statistics and metadata were compared from the StatsSpec and render metadata. Exports of four workloads in PNG, TIFF, PDF, SVG, and EPS at 300 dpi were validated with Pillow (dimensions, DPI tag, alpha, outer-ring clipping test), PyMuPDF (page size, extracted text against the expected title, axis labels, tick labels, and legend entries, embedded font types, rasterization for the clipping test), xml.etree (SVG size, text elements, absence of glyph outlines), and a Document Structuring Conventions parser for EPS; no PostScript interpreter was available, so EPS was validated structurally. Performance was measured on one workstation (Intel Xeon Silver 4208, 2.10 GHz, 32 logical CPUs) under Windows 11 (CPython 3.11.15) and under Linux in WSL2 on the same host (Ubuntu 22.04, CPython 3.11.8), with the two systems run one after the other and no thread limits set. Because performance does not depend on biological content, the inputs were synthetic tables of realistic shape generated with a recorded seed (numpy default_rng(20260904 + size)) and regenerated on demand: grouped long tables of 10^3^ to 5 × 10^5^ rows × 5 columns, DESeq2-shaped result tables of 10^4^ to 10^5^ rows × 8 columns, and Poisson count matrices of 100 × 12 to 50,000 × 100 features × samples; no real dataset was expanded or resampled, and no input larger than 5 × 10_5_ rows was tested. Load, analysis (Welch tests with Benjamini–Hochberg adjustment for the long table; log_10_ CPM, row z-scores, and PCA for the matrix), render, and export (PNG and PDF) were timed separately with one untimed warm-up followed by five timed repetitions (wall-clock and CPU time; peak resident-set increment sampled every 2 ms), then two repetitions under tracemalloc. The runtime in Fig. 4E is the median of the five per-repetition totals, with interquartile ranges in the benchmark tables deposited with the source code; peak memory is the tracemalloc Python-heap peak of the most demanding step (median of the two repetitions) and excludes process memory outside the Python allocator. Every plotted point, its input dimensions, replicate count, and origin are listed in the benchmark tables deposited with the source code. The technical tests were run on release v1.1.0 (commit 3da8563) on 14 September 2026 with all 38 registered plot types. Repeatability rendered each of the seven workloads and each registered plot type from its bundled example twenty times in one process on each system and compared every render with the first by structural signature, canonical SVG (volatile identifiers removed), and PNG pixels. Two defects found by these tests were fixed before the reported runs: the delimited loader’s float conversion, which now uses the round-trip converter, and the application of the style font stack to files saved outside the render context.

### Published-data figure recreation

For Fig. 5, candidate panels were sought in open-access articles from Nature-family, Science-family and Cell Press journals that deposit the values behind each figure panel as a Source Data worksheet or a supplementary table. Four literature searches yielded 43 candidate panels; ten were shortlisted and six selected by rules applied in order: the deposited table must hold exactly the numbers of one published panel, checked against the n, counts or statistics printed in its legend; the article must be CC BY so that the data and a reference image can be stored; no volcano plot and no statistical overlay, which are examined in Fig. 6; at least five plot families and no repeated journal; and legibility in a 58-mm cell without a panel-specific modification of the renderer. For each selected article the full-text XML and the supplementary files were retrieved from Europe PMC on 4 September 2026, the license element was read from the XML (all six articles are CC BY 4.0), and the raw workbook was stored unchanged with its SHA-256 checksum. Worksheets were exported to plain tables with pandas; the only operations were renaming of columns, reshaping of wide blocks to long form, and the transformations stated in the source legends. The six sources are: Gao et al. 2024^36^, Fig. 4c (hiPSC part), sheet “Fig. 4c”, 373 tRNA transcripts with the RPC1 ChIP–seq and tRNA-abundance fractions as deposited (the neuron columns belong to the other half of the panel and were not used); Ferretti et al. 2024^37^, Fig. 4a (left), sheet “Fig4a”, five treatment arms × five mice measured at up to 28 time points (525 values in the sheet; not every mouse at every time point), from which the renderer computes mean ± s.e.m. per arm and day, reproducing the sheet’s own mean and s.e.m. columns; Link et al. 2025^38^, Fig. 2h, sheet “Fig.2H_oncoPrint”, ten genes × 271 tumors with one alteration class per cell (736 altered cells), cells listing two alterations recoded as “Multiple alterations” and the 15 tumors without an alteration not drawn; Daugelaite et al. 2025^26^, Fig. 3d, sheet “Fig3d”, 40 samples × 38 transcription factors of per-factor row z-scores (mean 0 and s.d. 1 per factor verified), transposed to factors × samples, condition read from the sample identifier; Secchia et al. 2022^39^, Figure 1D, Table S1 (mmc2.xlsx), the 21,307 nuclei with published UMAP coordinates, which are the nuclei passing the authors’ quality filter; and Milotay et al. 2025^40^, Fig. 2d, sheet “Fig2d”, 75 patients with serostatus, months to death or censoring, and event indicator.

Each table was rendered by the registered renderer named in the figure legend through the same functions the interfaces call, with the column mapping recorded in the PlotSpec. Rows of the heatmap were kept in the deposited order, which is the published top-to-bottom order; because that order is not a leaf ordering of a Euclidean complete-linkage tree of the deposited z-scores, the rows were not re-clustered, whereas the samples were clustered with Euclidean distance and complete linkage and the tree drawn as in the paper. The Kaplan–Meier estimate, the regression fit and the mean ± s.e.m. bands were computed by the renderers. For every panel the plotted quantities were recomputed independently from the exported table with numpy, pandas, SciPy and the application’s log-rank routine and compared with the renderer metadata and with the values printed in the published panel or legend (sample sizes, R^2^, alteration frequencies, matrix dimensions and per-row standardization, population counts, numbers at risk at six-month intervals, survival fractions and the published log-rank P; 34 checks, all passed). The published panel images were compared visually with the recreations and are stored for review; they were not used as inputs. Reproduction was classified as exact analytical reproduction (original software rerun), scientific reproduction (same observations, transformation and plot form; drawing style differs), publication-grade recreation (same data, plot form changed) or new visualization; all six panels are scientific reproductions. Panels were rendered at 58-mm width with the manuscript typography (Arial; panel letters 14 pt), assembled by the application’s Figure Builder in a 3 × 2 grid with 600-dpi panel embedding and vector letters, and exported as PDF, SVG and PNG with the FigureSpec sidecar; the numerical content was frozen by checksum before and verified unchanged after the visual refinement.

### Independent R statistical validation

For Fig. 6A,B, every statistical test, multiple-testing correction, matrix transformation, quality-control metric, principal component analysis, hierarchical clustering, per-feature differential screen and renderer-derived statistic in the core was re-implemented independently in R 4.3.3^72^ (base R; survival 3.8-3^73^, car 3.1-3^74^, rstatix 0.7.2^75^, effectsize 1.0.1^76^, MASS 7.3-60^77^, dunn.test 1.3.6^78^, pROC 1.19^79^, limma 3.58.1^30^, edgeR 4.0.16^24^, DESeq2 1.42.0^29^) and run on 26 frozen synthetic datasets (seed 20250904), the 11 bundled example tables and the GEO GSE299655 RSEM count matrix^23^. The R scripts read only the input data and never any Python output; both sides write long-format result tables from independent code, and one comparison script merges them. Where a documented convention had to be matched (SciPy’s exact-versus-asymptotic rule for the Mann– Whitney U, Breslow ties in Cox regression, population z-scores), the R script applies the same convention and reports R’s default alongside as unscored context. Comparison classes were prespecified: exact (|Δ| ≤ 10^−12^), numerically equivalent (|Δ| ≤ 10^−10^ absolute or ≤ 10^−8^ relative), acceptable implementation difference (same method, documented cause), upstream input difference (input differs through the trimmed-mean trim-boundary tie handling), method mismatch (different model) and failure. The full matrix comprises 2,625 comparisons (Supplementary Fig. 2; statistics 1,039; multiple testing 1,176; transformations 88; quality-control metrics 112; PCA 15; clustering 31; per-feature screen 100; renderer-derived statistics 35; count-matrix characterization 29). Documented implementation differences are of four kinds: the generalized-linear-model standard error (R uses the weights of the final iteratively reweighted least-squares iteration and statsmodels the converged information matrix; relative difference ≤ 7.5 × 10^−8^ for Poisson and Gamma families and ≤ 3 × 10^−6^ for negative binomial with fixed dispersion), the Monte Carlo P value of r × c Fisher tests (≤ 1.9 × 10^−4^ from the exact value), trimmed-mean normalization factors at the trim boundary (ordinal versus average ranks; ≤ 4.4 × 10^−6^ on the RSEM matrix), which propagates to the log-CPM and voom rows classed as upstream differences, and the four-parameter logistic fit within the optimizer tolerance. Six numerical defects found by this validation (an unseeded r × c Fisher Monte Carlo, ROC and precision–recall curves that did not collapse tied scores, quantile normalization and voom that deviated from the limma definitions, and the sign of the Mann–Whitney effect size together with near-tie rounding in rank tests) were corrected and covered by regression tests before the reported run. Fig. 6B shows, for nine procedures, every compared test statistic, P value and effect size or estimate (143 quantities from 22 datasets; confidence-interval bounds are in the benchmark but not plotted) as the ratio of its absolute difference to its tolerance, max(10^−10^, 10^−8^ × max(|Python|, |R|)); exact zeros are drawn at 10⁻⁸ on the logarithmic axis.

### Published statistical analyses

For Fig. 6C,D,F, candidate analyses were sought in CC BY articles that deposit the source observations, not summary statistics, behind a panel, state the test, and print the P values, and whose test is one MakeMyFigure implements; 66 candidate analyses from 27 articles were recorded and three were selected, one two-group, one multi-group and one survival analysis, from journals not used in Fig. 5 where possible. Source Data workbooks were downloaded from the publisher (SHA-256 recorded) and the worksheet was exported verbatim and reshaped to one observation per row without changing any value. For Fig. 6C (Wu et al. 2025^42^, Fig. 4h; sheet “Fig. 4h”; 19 mice, n = 6, 6 and 7) the authors’ pairwise two-sided Welch *t*-tests^27^ without correction were run in MakeMyFigure (box plot with statistical annotation; StatsSpec test welch_t, all pairs) and in R as t.test(var.equal = FALSE) on the same observation table, with Hedges’ g as the effect size on both sides. For Fig. 6D (Wilton et al. 2023^43^, Fig. 6a; 63 participants, n = 13, 18 and 32) the authors’ Kruskal–Wallis test followed by Dunn’s test with Bonferroni correction over the three pairs was run in MakeMyFigure (StatsSpec test kruskal_wallis, post hoc dunn, correction bonferroni) and in R as kruskal.test with Dunn’s z computed from mean ranks with the tie correction and p.adjust(method = “bonferroni”). For Fig. 6F (Till et al. 2024^45^, Fig. 2A; sheet “Figure 2”; 56 patients in the sheet, of whom the 33 with circulating KRAS G12D and a recorded variant-allele-fraction group were retained, 17 at or below and 16 above the median, 27 deaths) the Kaplan–Meier estimate^41^ and the log-rank test^65^ were computed by the application and the Cox hazard ratio^66^ with Breslow ties by its regression module, and compared with survfit, survdiff (rho = 0) and coxph(ties = “breslow”) from the survival package. Agreement between MakeMyFigure and R was judged with the benchmark tolerances (10^−10^ absolute or 10⁻⁸ relative); agreement with the published values was judged at the precision printed in the article, and a printed threshold (“> 0.999”) was treated as a threshold. Every P value on the panels is formatted by the statistics engine from the stored StatsSpec.

### Software testing

The core is covered by 1,755 pytest tests in 99 files on the figure-package branch (commit 193f637), 1,709 of them outside the Qt interface module, including renderer tests for every plot type from its bundled example, statistics tests pinned to SciPy and statsmodels, regression tests for each defect found by the R validation and by the technical tests, preset round-trip tests for all plot types, Figure Builder geometry tests, loader and workbook tests, and interface tests for both frontends (the Qt tests run where PySide6 is installed). Tests run with the Agg backend.

### Figure preparation

All analytical panels in Figs. 3, 5 and 6 and panels D and G of Fig. 2 were rendered by make_my_figure_core using the same registry functions called by the graphical interfaces. The schematic panels A–F of Fig. 2 were drawn from the workflow’s JSON records, and the status matrices, bars and scaling plots of Fig. 4 were generated from the benchmark tables with matplotlib.

Figs. 5 and 6 were assembled using the application’s Figure Builder. Figs. 2–4 were composed from exported vector PDFs on a 183-mm column grid using PyMuPDF, except for the two composite examples within Fig. 3, which were generated by the Figure Builder. Figure Builder composites used two-or three-column grids at 183-mm width, with panel content embedded at 600–1,200 dpi and panel letters retained as vector text; a FigureSpec sidecar was written with each composite. Panel letters were 14-point Arial. Fig. 1 was drawn with matplotlib primitives and edited by the authors in Adobe Illustrator. Typography was checked on the rendered PDFs, and the numerical content of every panel was frozen by SHA-256 checksums of the input tables and specifications before visual refinement and verified unchanged afterward.

## Data availability

All datasets are public. Source Data workbooks and supplementary tables of the six Fig. 5 articles^26,36–40^ and the four Fig. 6 articles^42–45^ are distributed with the articles under CC BY 4.0; the GSE299655 count matrix and sample records are at the Gene Expression Omnibus^23^; the penguin data are from palmerpenguins (CC0)^31^; Gapminder 2007 is from gapminder.org (CC BY 4.0)^32^; the Liu et al. 2018, Guo et al. 2019, and Kume et al. 2024 tables are from the articles’ source data (CC BY 4.0)^33–35^; the Osipovich et al. 2023 table is S2 Table of that article (CC BY 4.0)^71^. The exported tables used for each panel, the PlotSpec and StatsSpec sidecars, the per-panel audit tables, and the R-validation benchmark (inputs, R scripts, Python exports, comparison tables) are deposited with the source code.

## Code availability

MakeMyFigure is released under the MIT license at https://github.com/surPoudel/make-my-figure, together with the User Manual and Quick Start guide (Supplementary Notes 1 and 2), the benchmark folders (benchmarks/r_validation and the technical-validation scripts), and the scripts that produced Figs. 2–6.

## Acknowledgements

This work was supported by the Anthropic AI for Science program, which provided API credits used for software development, code review, and documentation of MakeMyFigure. The content is solely the responsibility of the authors and does not necessarily represent the official views of the funders. The funders had no role in the design of the software, the analyses, the decision to publish or the preparation of the manuscript. Claude (Anthropic) was used under author supervision to assist with code generation, refactoring and copy-editing; the authors reviewed and verified all code and text and take full responsibility for the content.

## Author contributions

S.P. conceived the project, applied for and obtained funding from the Anthropic AI for Science program, and designed and developed the software from its inception to the released versions. H.K.S. tested successive versions of the software, provided feedback and helped write the manuscript. J.C.C. reviewed the manuscript and provided critical feedback. F.D. tested the software and reviewed the manuscript. D.R.G. supported S.P. in preparing the funding application, provided feedback and suggestions throughout the development of the software and reviewed the manuscript. All authors read and approved the final manuscript.

